# Measuring the accuracy of gridded human population density surfaces: a case study in Bioko Island, Equatorial Guinea

**DOI:** 10.1101/2020.06.18.160101

**Authors:** Brendan F Fries, Carlos A Guerra, Guillermo A García, Sean L Wu, Jordan M Smith, Jeremias Nzamio Mba Oyono, Olivier T Donfack, José Osá Osá Nfumu, Simon I Hay, David L Smith, Andrew J Dolgert

**Affiliations:** South and Central Africa ICEMR, Johns Hopkins Bloomberg School of Public Health, Baltimore, MD, USA; Spatial Science for Public Health Center, Johns Hopkins Bloomberg School of Public Health, Baltimore, MD, USA; Medical Care Development International, Silver Spring, MD, USA; Divisions of Biostatistics & Epidemiology, University of California, Berkeley, CA, USA; Medical Care Development International, Malabo, Equatorial Guinea; Ministry of Health and Social Welfare, Malabo, Equatorial Guinea; Department of Health Metrics Sciences, School of Medicine, University of Washington, Seattle, WA, USA; Institute for Health Metrics and Evaluation, University of Washington, Seattle, WA, USA

**Keywords:** population health, demographics, epidemiology, mapping

## Abstract

Geospatial datasets of population are becoming more common in models used for health policy. Publicly-available maps of human population in sub-Saharan Africa make a consistent picture from inconsistent census data, and the techniques they use to impute data makes each population map unique. Each mapping model explains its methods, but it can be difficult to know which map is appropriate for which policy work. Gold-standard census datasets, where available, are a unique opportunity to characterize maps by comparing them with truth. We use census data from Bioko Island, in Equatorial Guinea, to evaluate LandScan (LS), WorldPop (WP), and the High-Resolution Settlement Layer (HRSL). Each layer is compared to the gold-standard using statistical measures to evaluate distribution, error, and bias. We investigated how map choice affects burden estimates from a malaria prevalence model. Specific population layers were able to match the gold-standard distribution at different population densities. LandScan was able to most accurately capture highly urban distribution, HRSL matched best at all other lower population densities and WorldPop performed poorly everywhere. Correctly capturing empty pixels is key, and smaller pixel sizes (100 m vs 1 km) improve this. Normalizing areas based on known district populations increased performance. The use of differing population layers in a malaria model showed a disparity in results around transition points between endemicity levels. The metrics in this paper, some of them novel in this context, characterize how these population maps differ from the gold standard census and from each other. We show that the metrics help understand the performance of a population map within a malaria model. The closest match to the census data would combine LandScan within urban areas and the HRSL for rural areas. Researchers should prefer particular maps if health calculations have a strong dependency on knowing where people are not, or if it is important to categorize variation in density within a city.

## Introduction

With the implementation of national malaria elimination campaigns in many countries across the globe, planning, monitoring, and evaluation of malaria interventions have become more critical than ever (1). Advances in mapping and modeling of disease risk and spread have accelerated since the turn of the century, accomplished through increases in GIS and satellite imagery and survey data (2, 3). These advances have led to the creation of publicly available global population datasets that have often been used to inform many public health studies in areas without first-rate census data, as in most of the developing world, where much of the infectious disease burden resides. The inherent uncertainty in models and estimates of infectious disease burden is usually recognized while the fundamental uncertainty in the denominator of such disease estimates, the human population data layer, is commonly assumed to be completely correct.

The premise of precision public health is that evidence can be used to improve the efficiency and effectiveness of interventions to benefit those most in need (4, 5). Data describing the geographical distribution of humans—accurate human population density maps—are among the most important components of evidence, along with observational or intervention studies, as they describe the needs for resource provisioning and the denominators for the analyses used to identify populations at risk. Risk factors, health catchment populations, access to resources, and operational constraints on provisioning resources are intrinsically related to population density and geographical location. In the context of malaria, population surfaces are also increasingly used as covariates for geostatistical models for mapping prevalence, incidence and other metrics (6–9). Information about the geographical location of human households is required to weigh resource allocation decisions to identify and serve populations in need (10, 11). Making effective policy thus requires having accurate maps of human populations (10). The last two decades have seen significant advances in mapping technologies, and the publication of several gridded population surfaces using different modeling approaches (11).

The basic notion of accuracy for population maps is quite intuitive. Information in maps is complicated, however, and while a perfect map is accurate in every way, an inaccurate map can be inaccurate in many different ways. An open question is how to measure the accuracy of maps for different purposes. Two questions for precision public health are how to measure the accuracy of these maps, and how to set standards for accuracy for various purposes. Maps need not be perfect, but they should be suited to the task at hand.

Here, we measure the accuracy of published overlapping maps of human population density using a high quality map of one locale as a gold standard (12). We use several metrics to evaluate these maps, including a new goodness-of-fit metric and a new application of accuracy profiles. We evaluate their suitability in the context of malaria control and elimination policy on Bioko Island. The Bioko Island Malaria Elimination Project (BIMEP) has developed highly detailed and constantly updated housing cartography as a basis for distributing interventions, monitoring impact and implementing surveillance (12). Accompanying this housing database is a recent population census that allocates inhabitants to their households. We use these BIMEP population data as the gold standard against which we evaluate several publicly available gridded population surfaces. We also discuss the functional consequences of accuracy by using these surfaces to develop maps of *Plasmodium falciparum* parasite rate (*Pf*PR) using well-documented methods, where the population density surface is used both as a covariate and as the population weight on the *Pf*PR surface.

## Methods

### Study area

Bioko is the largest island of Equatorial Guinea, at 2017km^2^. It is located approximately 40km off the coast of Cameroon, in the Bight of Bonny. Malabo, the main urban centre and country capital, concentrates around 85 % of the human population of the island. Administratively, Bioko is divided into two provinces (second administrative division) and four districts (third administrative division; Figure 1).

**Fig. 1.**
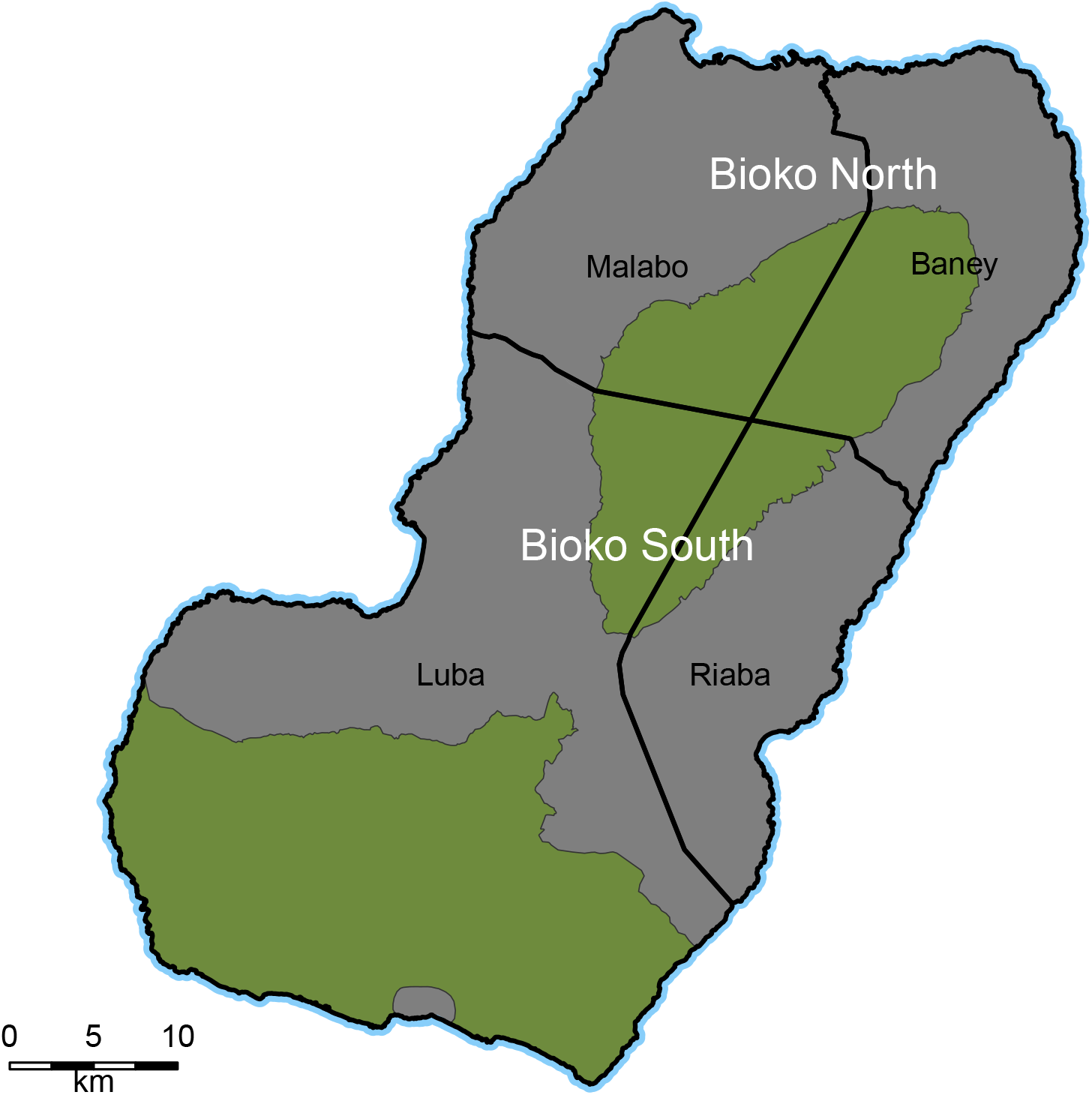
Second and third administrative divisions on Bioko Island. The thick black lines demarcate the four districts. Malabo and Baney make Bioko North and Luba and Riaba, Bioko South. Green areas are uninhabited nature reserves.

### Population data

#### BIMEP health census data

This health population census was part of a bed-nets mass distribution campaign in 2018 (13). People present during the campaign were counted and registered to their house mapping code for geographical reference (12). Each house is GPS-located on the island. The census underestimates the total household count by approximately 12 %, due to BIMEP census workers only being able to reach approximately 88 % of them during the bed net intervention campaign. The underestimate in the actual population is likely different, however, due to the heterogeneous distribution of intra-household population counts. Despite this discrepancy, the 2018 BIMEP population census represents the most accurate and most up-to-date rendering of population distribution on Bioko Island. This census followed a similar population count in 2015 during the preceding bednets distribution campaign, so the data were validated against this previous effort and are used here as the gold standard for analyses.

#### Gridded population data

We selected all population maps that are publicly available and nearly complete across Africa: WorldPop (WP) 2015 (14), LandScan (LS) 2017 (15) and the High Resolution Settlements Layer (HRSL) (16). WP utilizes a mixed approach for mapping populations that includes areal weighting of census data and dasymetric modelling, a type of thematic geospatial map that incorporates ancillary remotely-sensed and geospatial data (14, 17). The WP input population for Equatorial Guinea corresponds to third administrative level (district) census data dated 2001, and we used the layer that is adjusted to UN estimates of total population. LS uses census data and dasymetric mapping that incorporates multiple data layers including land cover, roads, slope and human settlements (15). The HRSL uses machine learning algorithms to map buildings at very high spatial resolution (1 arc second, or approximately 30 m) (16). All buildings identified in this layer are then proportionally allocated human population from second administrative level census data. All three datasets are on grids in latitude and longitude. At Bioko’s latitude, the median side length of a grid square varies from 30.8 m for HRSL, to 92.4 m for WP and 924 m for LS.

We needed to compute a density per square kilometer in order to measure urban fraction. While the gridded population surfaces can be treated as density per pixel, we generated separate population density surfaces by taking the sum of all people within a radius of 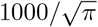 meters of the grid square centroid. This algorithm assumes the density within each pixel is constant.

When there is direct comparison between the house-level BIMEP data and a gridded dataset, we aggregated the BIMEP data to the same grid as the dataset. This way, each map is compared without added interpolation from alignment to a common grid. When algorithms called for BIMEP data to be gridded, we used the finest grid, that of the HRSL. Where the size of a grid cell might affect comparison, we aggregated the HRSL 32-fold and LS 11-fold, in order to construct nearly 1 km grids. We refer to these as 1 km maps.

It seems possible that the exact alignment of the raster grid over the island might affect metrics. In order to investigate this, we rasterized the BIMEP data hundreds of times to grids of the same resolution, but shifted slightly in latitude or longitude. We recalculated the basic metrics in Table 1 for each shift, and it is the standard deviation of the resulting distribution of metrics that appears as plus-minus values in that table. Of these metrics, the maximum population is most sensitive to grid placement, especially for the coarsest grid choice.

**Table 1.** Basic summary statistics for all the maps. The plus-minus values on the BIMEP columns reflect how much the exact grid alignment, in latitude and longitude, matters. We shifted the grid a hundred times in either direction and recalculated metrics. Only the maximum population density is sensitive to the exact location of pixels.

|  | BIMEP | HRSL | LS | WP | BIMEP | HRSL | WP |
| --- | --- | --- | --- | --- | --- | --- | --- |
| Grid Size | 985.3 m | 985.3 m | 923.7 m | 1016.0 m | 30.8 m | 30.8 m | 92.4 m |
| Total | 239056 | 231210 | 218044 | 204820 | 239056 | 231210 | 204820 |
| Max. Pop. Dens. | 17130±1200 | 6085 | 34344 | 1018 | 22212±125 | 7525 | 1017 |
| % Empty | 88.61±0.2 | 86.32 | 56.29 | 0.6536 | 99.13±0.001 | 98.92 | 1.137 |
| % Urban | 2.007±0.1 | 3.175 | 2.075 | 0.2011 | 2.029±0.1 | 3.283 | 0.1588 |
| Pareto Number | 3.875±0.001 | 5.696 | 5.275 | 24.13 | 3.904±0.000 | 5.643 | 24.7 |

#### Goodness of Fit Ratios

A commonly used distance metric to measure statistical models’ goodness of fit is the sum of squared errors (10); this metric is related to variance (average distance of a random variable from its mean) by noting that the mean sum of squared errors is equal to bias plus variance. We propose a new measure of the utility of a map is whether a measure of goodness of fit based on the sum of squared errors is better than the variance, which is the goodness of fit for a “null” map that assigned to each pixel the average population density. We call this a goodness-of-fit ratio (GOFR).

A GOFR of 0 indicates a perfect fit. A GOFR greater than one indicates the fit was worse than the null. A GOFR of 1 indicates that the map is as good as a null map. We apply GOFR both to the whole of Bioko and separately to each second administrative level.

Let *H*(*x*) denote the true population density at pixel *x*. Denote the observed map value with 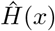. For any subset of map pixels, the GOFR is the ratio of mean squared error per pixel to variance of the expected. Using *n* for the number of pixels in a region, the GOFR for that region is

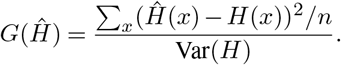

We can also compute a normalized by district GOFR to compare relative population densities, to remove any effect of having different total population sizes. Let *h*(*x*) = *H*(*x*)/∑_*y*_*H*(*y*). The normalized version is

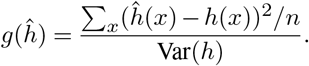

If *G* = 0 then the candidate map is perfect over the region. If *G* < 1, then the map improves the goodness of fit over the constant value, and if *G* ≥ 1, the goodness of fit is worse than just using the average population density.

#### Urban Fraction

Countries measure urban fraction in ways that are germane to their needs for planning and assessment, so the measure can include observations of human movement patterns and availability of resources. Because we are looking only at the maps, the urban fraction here is the percentage of pixels for which the density within a square kilometer was greater than a thousand people. It may be that Equatorial Guinea uses a cutoff of 1500 people per square kilometer (18), but the relative information in these maps is the same for either choice of cutoff.

#### Pareto Number

The Pareto number is a single value that characterizes the tendency of population to aggregate. A smaller value indicates more aggregation. For instance, if 95 % of the population is in 5 % of the pixels, the Pareto number would be 5. We find the Pareto number by sorting pixels in increasing population size. The Pareto number is the index, normalized to 100, of the pixel for which the fraction of pixels that are larger equals the fraction of total population in pixels that are smaller.

#### Accuracy Profiles

We used binary classification statistics to construct an accuracy profile for the 1 km maps (Table 2). Binarization of population quantity numbers was done to allow comparison of accuracy statistics across multiple population surfaces. For a threshold population density, *τ*, each pixel in a map is classified as being either above or below the threshold. Using BIMEP as the gold standard, we assessed the accuracy of the other maps against it and each other. We computed: true positives (TP), the proportion above the threshold in both maps; true negatives (TN), the proportion below the threshold in both maps; false negatives (FN), the proportion above in BIMEP but below in the other map; and false positives (FP), the proportion below the threshold in BIMEP but above in the other map. We define accuracy as the proportion correct (i.e. (*TP* + *TN*)/(*TP* + *TN* + *FP* + *FN*)); recall or sensitivity as the proportion above a threshold in the gold standard that were correctly assigned: *TP/(TP + FN*); and precision or positive predictive value as the proportion above the threshold in the alternative map that were correctly assigned: *TP*/(*TP* + *FP*). Each threshold value on population density, *τ*, gives different measures of accuracy, recall, and precision. We also computed accuracy metrics for classification of the landscape into population density categories: empty (strictly equal to zero), and for breakpoints at 1, 50, 250, and 1,000 people per km^2^ with 1,000 and up classified as urban areas.

**Table 2.** Let a threshold, *τ*, define a categorization of population density. In a gold standard map, *G*, a pixel is in the category if it is above the threshold: *x* ∈ *G_τ_* if and only if *x > τ*. Otherwise, 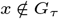. Similarly, the categorization is applied to a candidate map, *M*. Pixels are classified as true positives (TP), true negatives (TN), false negatives (FN), and false positives (FP) as described in the table. Accuracy profiles are plotted in Figure 6.

| $x \geq \tau$ | $x \in M_\tau$ | $x \notin M_\tau$ |
| --- | --- | --- |
| $x \in G_\tau$ | TP | FN |
| $x \notin G_\tau$ | FP | TN |

#### *Pf*PR mapping

We estimated the prevalence of malaria parasites, a *Pf*PR surface, for Bioko Island using the same set of covariates, replacing only the population surfaces one at a time. Data and methods described elsewhere (8, 9). The population surfaces were then used to construct density values to assign as population weights for use in the calculation of *Pf*PR. The response data corresponded to *Pf*PR at household-level spanning the period 2015–2018. We ran this exercise for each of the 1×1 km population grids since environmental covariates were not available for Bioko at finer spatial resolution. We also estimated relative populations at risk using each population surface and expressed them as cumulative distribution and probability density functions.

## Results

### Population distribution

Much of the habitable land area on Bioko Island is sparsely inhabited and the bulk of the population is concentrated in the North, within and nearby Malabo. The rest of the population is distributed in pockets, mostly rural, along the East and West coasts. There are two large, uninhabited nature reserves in the North and South of the island (Fig. 1). Fig. 2 and 3 illustrate the population distribution according to each of the four surfaces at 1×1 km and 100×100 m, respectively. In the BIMEP surface (Fig. 2A), the population is highly concentrated around the center of Malabo, with areas housing as many as 17,130 people per square kilometer. Fig. 3A illustrates a highly heterogeneous human population distribution in the center of Malabo at 100×100 m pixels. Given the small size of Bioko and aggregation of its population, it’s a success for the metrics to present a consistent picture of map performance.

**Fig. 2.**
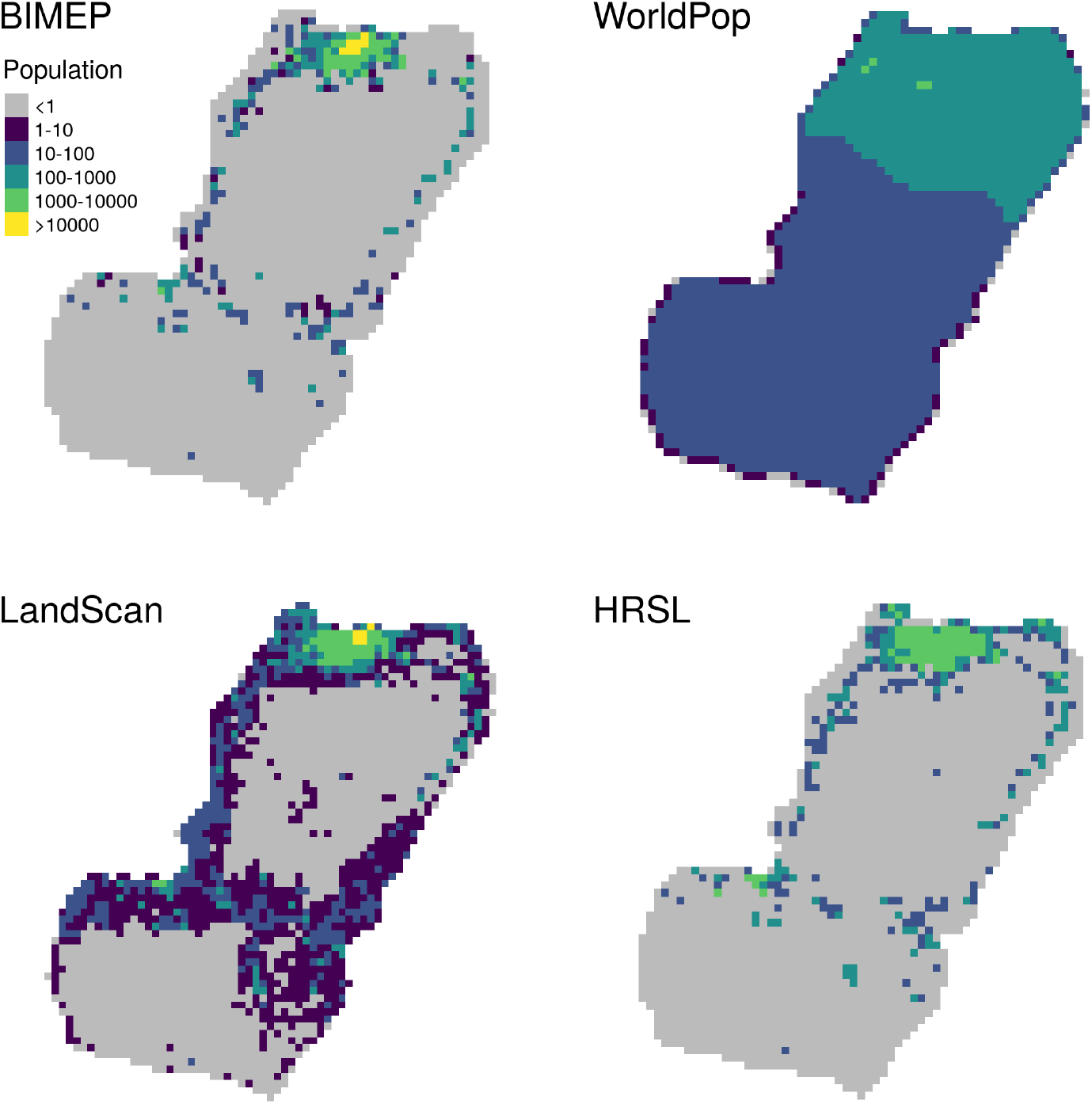
Bioko population rendered at 1×1 km resolution. A. BIMEP; B. WP; C. LS; D. HRSL. Grey pixels represent uninhabited areas (population = 0).

**Fig. 3.**
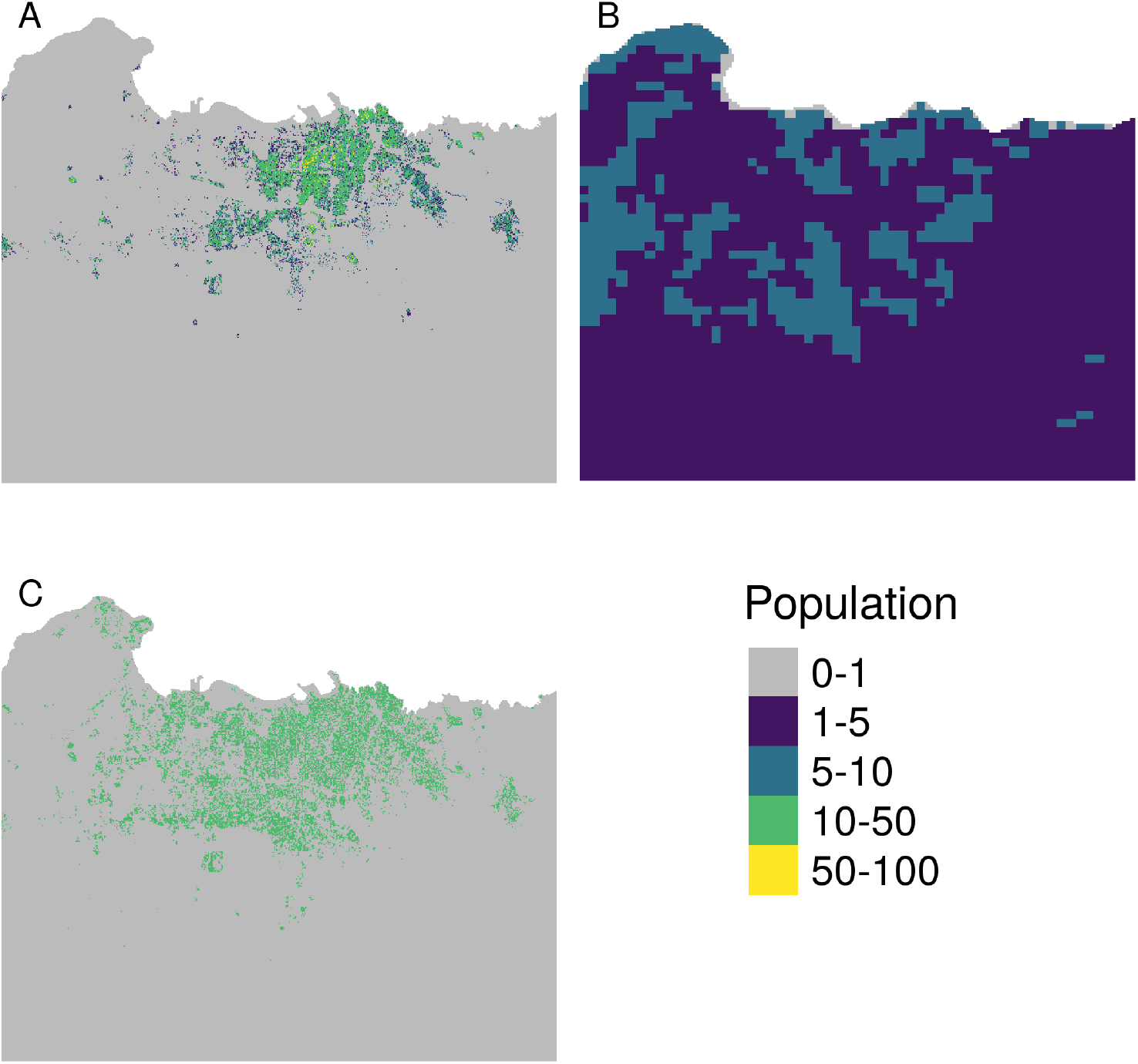
Bioko population rendered at 100×100 m spatial resolution. The maps are zoomed into the Malabo area as visualization of a highly populated area for comparison. A. BIMEP; B. WP; C. HRSL. Grey pixels represent uninhabited areas.

The population according to WP distinguishes average density between Bioko North and Bioko South but diverges little from mean values, including within the nature reserves (Fig. 2B). The 100×100 m WP surface does show increased densities around the Malabo area, but they intersperse with low populations in the urban city center (Fig. 3B).

The 1 km LS population surface, while correctly identifying the general shape of settlement around the Malabo area, fails to predict the extreme abundance of zero population pixels in the rest of the island. Within Malabo, the LS model severely over-predicts extreme population aggregation, estimating 34.6 % of the total population is concentrated amongst two square kilometers with population density as high as 34,344 people/sq km (Fig. 2C).

The HRSL surface renders a more accurate population distribution overall, particularly in rural areas (Fig. 2D). It fails to provide an accurate picture of urban Malabo, however, where the population appears more evenly distributed than the gold standard, with a maximum population density of 6,085 per square kilometer. This pattern is also manifest in the 100×100 m HRSL surface, representative of an overly uniform population distribution across Malabo (Fig. 3C).

### Per-pixel Scatter Plot

The biases and exactness of the algorithms used to generate the maps are more obvious in the scatter plots (Fig. 4). In particular, the raw plots for the 30×30 m and 100×100 m maps for HRSL and WP show distinct horizontal striping patterns (Fig. 4E,F). The distinct horizontal stripes observed in these plots indicate that for a large range of actual (BIMEP) population densities, WP and, to a lesser extent, HRSL predicted constant density; this is a visual indication of model inability to fully characterize spatial variation in population density. The HRSL’s estimated map is produced by an algorithm which, in this case, identifies a maximum of 20 households in each pixel, and allocates the total population evenly among households, producing here an integer multiple of 10.144 individuals to each household. In WP, each grid square is assigned to one of seven distinct population density values. These patterns are obscured in population density estimates or in aggregating data up to 1 km grid cells. The adjusted *R*^2^ values for the per-km maps are higher than for their respective estimated population density, which are higher than for the population size as well.

**Fig. 4.**
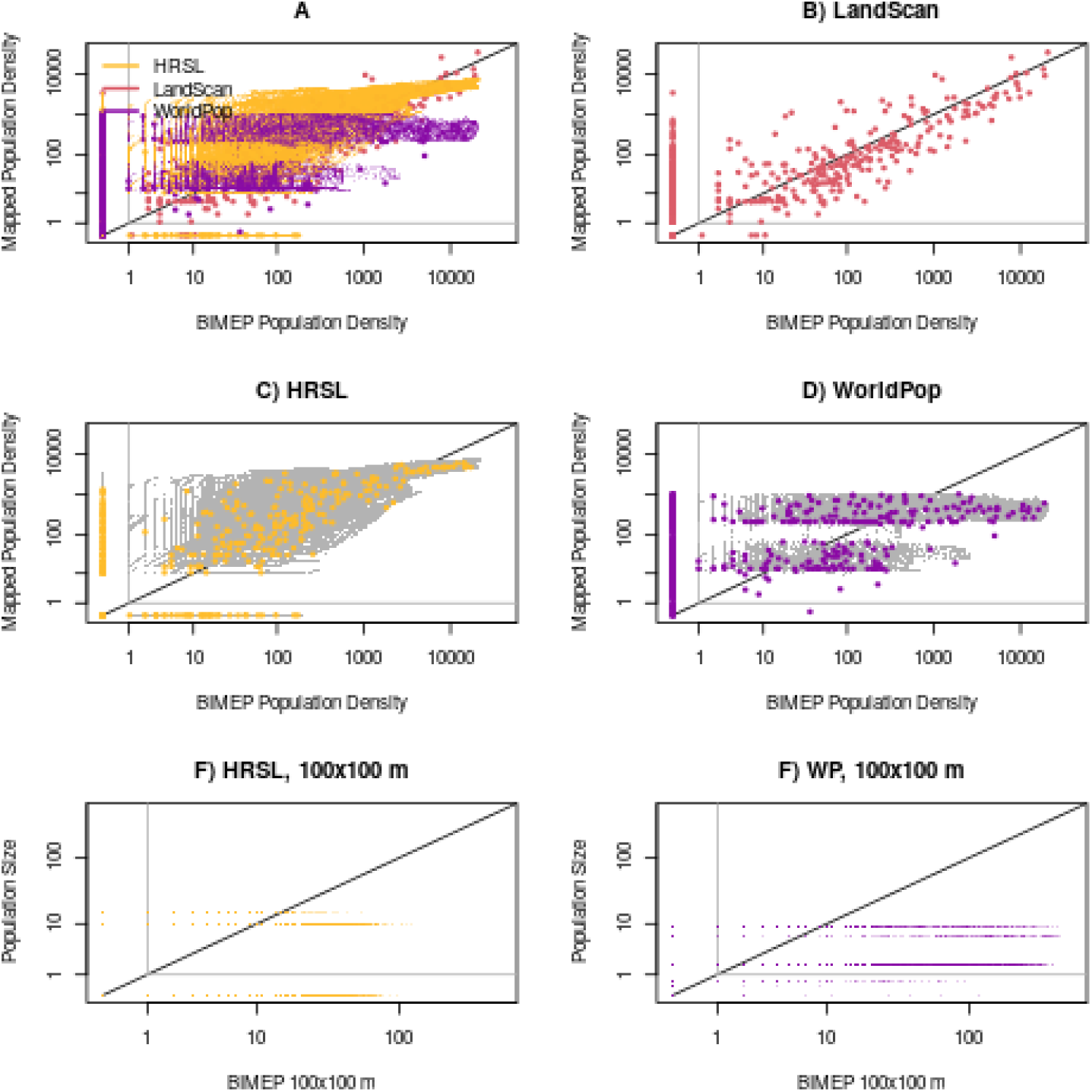
Scatter plots reveal the accuracy and biases of the algorithms used to generate the maps. The one-to-one line is plotted in black and the grey vertical and horizontal lines are plotted at a population density or size of 1. Figure A has a color legend for all panels. A-D) Scatter plots of the population densities from all the different population surfaces plotted against the gold standard of BIMEP. E-F) The population size for the 100×100 m maps. A) All 100 *m* and 1 *km* maps are plotted on the same axis. The 100 *m* pixels are smaller but follow the same patterns as the corresponding 1 *km* maps. The visual impression is dominated by the 100 *m* maps (because there 10,000 times more pixels), so we have also plotted the 1; *km* maps, if relevant, with the 100 *m* map as a grey background. B) Landscan at 1 km. Adjusted *R^2^* was 43%; C) HRSL at 1 *km* (yellow) and 100 *m* (grey). Adjusted *R*^2^ was 65% for the 1 *km* map and 56% for the 100 *m* map; D) WorldPop at 1km (yellow) and 100 *m* (grey). Adjusted *R*^2^ was 5% for the the 1 *km* map and 3% for the 100 *m* map. E) Population size in the BIMEP vs. HRSL maps at 100 *m*; adjusted *R*^2^ was 34%. F) Population size in the BIMEP vs. WorldPOP maps at 100 *m*. Adjusted *R*^2^ was 2%.

### Cumulative Distribution by Area

Some of the same patterns are evident in the empirical cumulative distribution functions (eCDFs) and smoothed density plots showing population density and its distribution by land area (Fig. 5), which highlight the large fraction of empty space in most of the maps (Fig. 5A). In the BIMEP maps, the fraction of empty pixels was 98 % for the 100m map and 88.4 % for the 1km map; both the HRSL and LS were similar (Fig. 4B). WorldPop, by way of contrast, reported a positive population density for almost every pixel. We have also plotted the eCDFs of population density vs. the proportion of the human population living at that density, and also the empirical probability distribution functions (Fig. 5B,C). These maps highlight important differences, such as maximum population density: 34,344 per km^2^ for the LS map, 17,130 for the BIMEP map, 6,085 for the HRSL map, and 1018 for WP (Table 1).

**Fig. 5.**
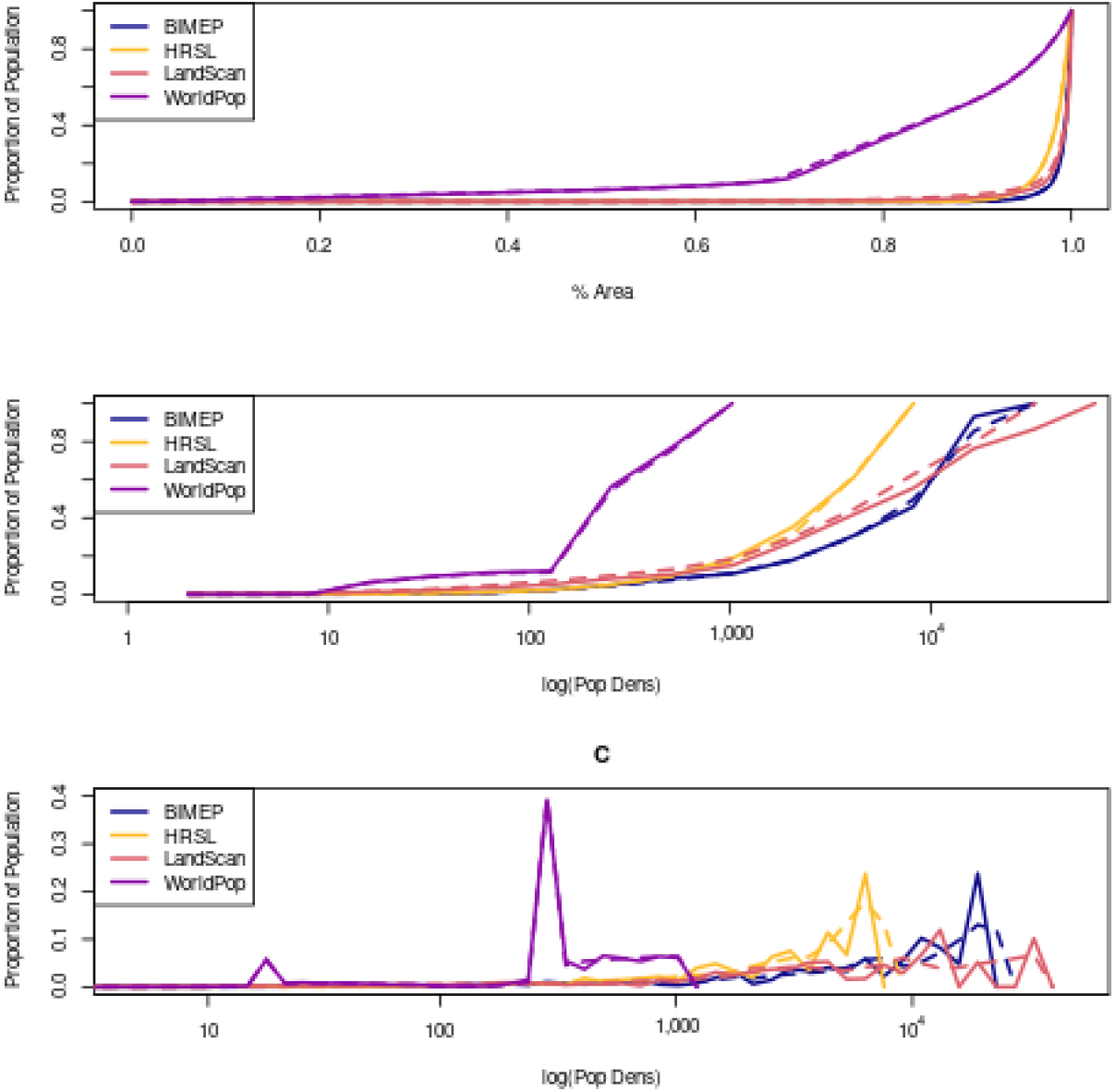
A comparison of the distributions by land area and population density. Solid lines are 1km maps, and dashed lines are 100m maps. A) To show how the population is distributed, we plotted the empirical cumulative distribution functions (eCDFs) of population density by land area; B) To show how the population is aggregated, we plotted the eCDFs by log population density. C) The population density binned by powers of 1.2.

### Goodness of Fit Ratios

We applied the goodness of fit ratio (GOFR) to all population maps at the 1 km resolution for comparison. The HRSL was an improvement over the average population density for all of Bioko Island and for each one of the third administrative levels (Table 3). The LS map was an improvement for all but the Luba district. The WP map was remarkably close to an average population density map. The GOFR values for Luba and Riaba were larger, in general. A small registration error in a map could result in a large GOFR because their populations are small (5500 in Luba, 2300 in Riaba) and highly aggregated.

**Table 3.** This compares the goodness-of-fit ratio across the three maps, aggregating HRSL and WorldPop to 1 km resolution to match LandScan. The HRSL is an improvement overall and, after normalizing, provides a good fit in each one of the districts. Normalization discounts the effect of uniform changes in population size. WorldPop is approximately as informative as the “null” map.

|  | GOFR |  |  | Normalized GOFR |  |  |
| --- | --- | --- | --- | --- | --- | --- |
|  | HRSL | LS | WP | HRSL | LS | WP |
| Bioko Island | 0.383 | 0.472 | 0.956 | 0.374 | 0.545 | 0.953 |
| Baney | 0.150 | 0.374 | 1.028 | 0.140 | 0.136 | 0.991 |
| Luba | 2.266 | 5.484 | 1.012 | 0.266 | 1.658 | 0.992 |
| Malabo | 0.412 | 0.487 | 0.987 | 0.377 | 0.570 | 0.974 |
| Riaba | 2.381 | 0.824 | 1.157 | 0.825 | 0.458 | 0.996 |

### Accuracy Profiles

The accuracy profile shows measures of accuracy, recall, and precision as binary classification statistics for a mesh on *τ* for values spanning the range of the gold standard (Fig. 6). The BIMEP and HRSL maps were the most similar across all three binary metrics. LS and WP had a much higher fraction of pixels in the lowest population category (Fig. 6A). Overall, the HRSL tended to be the most accurate and with the best recall for population densities up to 5,000 per km^2^, which was the upper limit of population densities in that map (Fig. 6B,C). The precision of the LS map was highest from 250 people per km^2^ up to around 2,000 people per km^2^ (Fig. 6D).

**Fig. 6.**
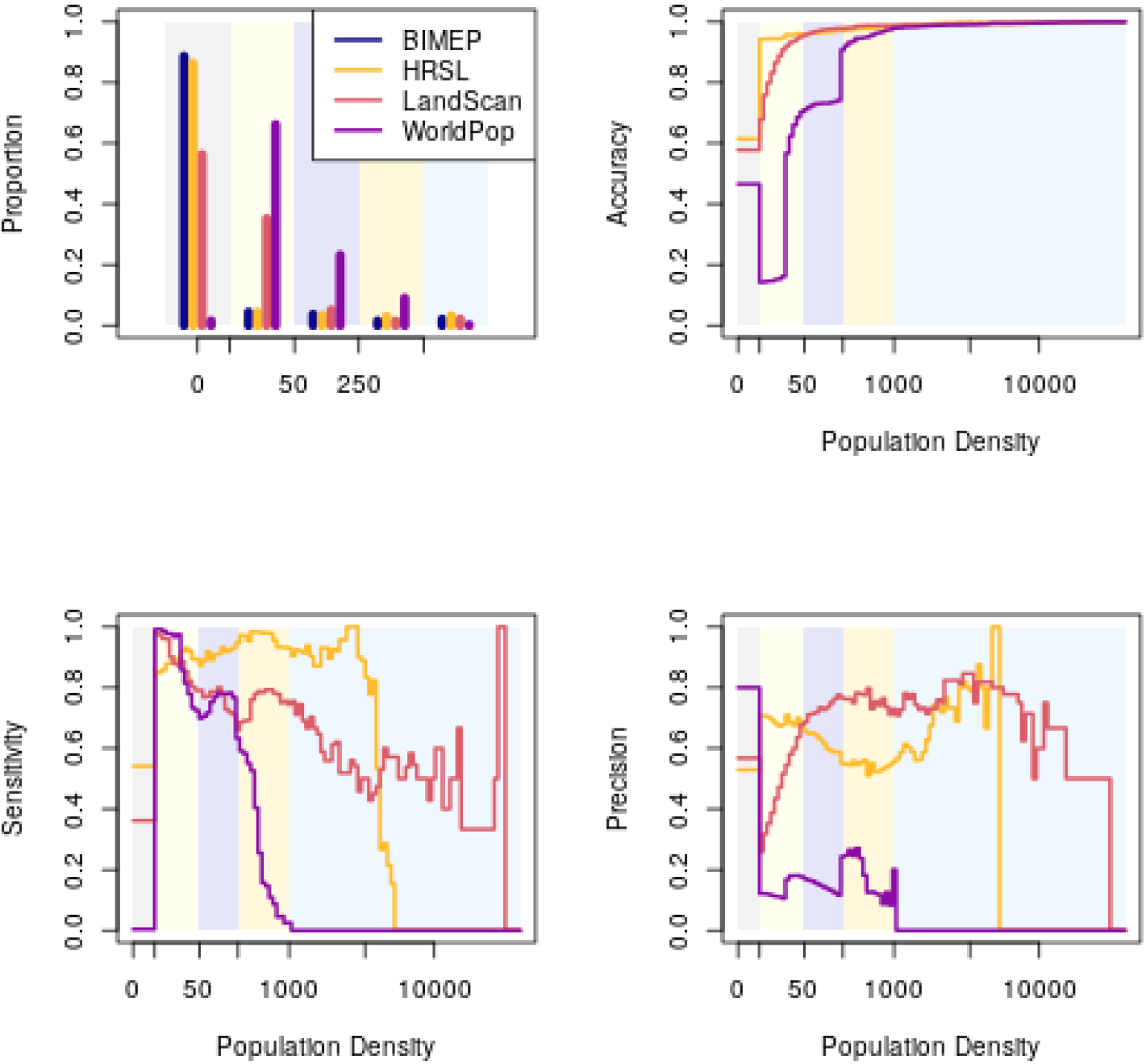
A) The proportion of the population in density categories defined by breakpoints of 1, 50, 250, and 1,000 people. B) The accuracy profile; C) The recall profile; D) The precision profile.

The HRSL map identified 96 % of the empty pixels and 90 % of the urban pixels correctly (i.e. by recall or sensitivity), while LS identified 98 % of the urban pixels correctly. Notably, the accuracy metrics are dominated by true negatives, since there are so many empty cells (Table 4).

**Table 4.** The accuracy, recall, and precision for the population classifications shown in the header and illustrated in Figure 6.

| $H$ | (0, 1) | [1, 50) | [50, 250) | [250, 1000) | [1000, ∞) |
| --- | --- | --- | --- | --- | --- |
| **Accuracy** |  |  |  |  |  |
| HRSL | 0.9430 | 0.9410 | 0.9600 | 0.9700 | 0.9850 |
| LS | 0.6720 | 0.6740 | 0.9530 | 0.9750 | 0.9890 |
| WP | 0.1290 | 0.3440 | 0.7540 | 0.9120 | 0.9800 |
| **Recall** |  |  |  |  |  |
| HRSL | 0.9550 | 0.3120 | 0.4160 | 0.4520 | 0.9070 |
| LS | 0.6320 | 0.7500 | 0.5380 | 0.2270 | 0.7450 |
| WP | 0.0124 | 0.4880 | 0.3010 | 0.5150 | 0.0256 |
| **Precision** |  |  |  |  |  |
| HRSL | 0.9800 | 0.3150 | 0.4440 | 0.2300 | 0.5740 |
| LS | 0.9970 | 0.0869 | 0.3770 | 0.3030 | 0.7290 |
| WP | 0.9570 | 0.0312 | 0.0542 | 0.0955 | 0.2500 |

HRSL had the highest precision (i.e. positive predictive value) for “empty”: if a pixel was reported empty in LS, it was empty in the BIMEP map 99 % of the time. The HRSL was a close second at 98 % PPV. The PPV values for urban classification were lower: LS was the highest at 77 %, while HRSL had 56 % (Table 4).

### Malaria Mapping

In our analysis, human population density was only weakly correlated with *Pf*PR on Bioko Island, so the resulting *PfPR* maps were virtually indistinguishable. The main difference was how the distribution of people affected calculation of average *Pf*PR: 11.3% for the BIMEP map, 12.3% for the HRSL, 13.4% for WP, and 10.6% for LS (Fig. 7A). Calculating the fraction of the population at greatest risk, with *Pf*PR above 20%, was sensitive to the map’s ability to identify urban areas, leading to 2.5% in BIMEP, 2.6% in LS, 6.2% in the HRSL, and 12.4% in WP (Fig. 7B).

**Fig. 7.**
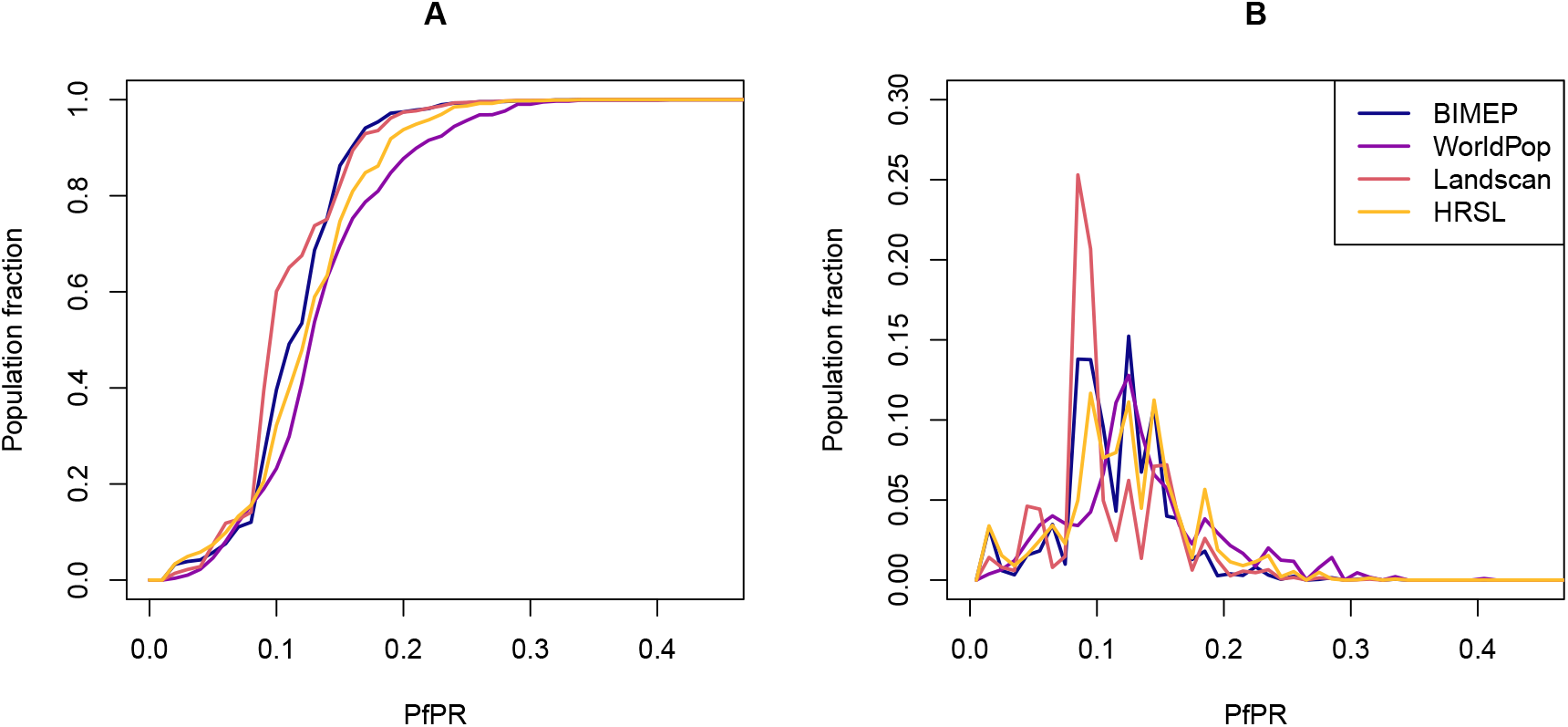
Fraction of the population according to *Pf*PR, expressed as cumulative distribution (A) and probability density functions (B).

## Discussion

The individual metrics, described above, together provide insight into optimal use and limitation of these datasets, both for malaria and wider public health applications. Comparing with a gold standard dataset gives us some sense of the effects each approach’s assumptions and use of ancillary data.

### WorldPop

The WorldPop results consistently under-reported population levels in urban areas (Fig. 6). In Table 1, WP has the lowest population totals and lowest max population density, this is likely due to WP maxing out population at 1000 people per km^2^, which is a low estimation for a primarily urban population like we see in Bioko Island where a majority of the population lives in Malabo. The average population density of Malabo is around 14,000 per km^2^ (9) which results in a significant under count, and this effect is increased when the resolution is increased to 100 m. On the opposite end of the population spectrum, WP severely overestimates population densities everywhere else, with the lowest number of empty pixels (Fig. 5). This has been seen before in other African countries and may be a product of the random-forest model approach (14). WP is the only surface to show people living in the large nature reserves in the North and South of the island (Figure 1). Both underestimation of high population areas and overestimation of low population areas may be the result of the WP population model being anchored by official census data at the second administrative division (14). Heavy reliance on the mean of census data could be problematic for countries where census data are reported only at higher administrative levels.

### LandScan

The LandScan global gridded population surface was the best at characterizing high-density urban population distribution but was the only of the three to overestimate population density compared to BIMEP (Table 1). The LS 1 km grid layer had the highest accuracy and precision in Malabo and high density (greater than 1000 people per pixel) areas (Table 3). If the two highest density pixels were removed, the LS layer’s GOFR scores improved significantly. Similarly, LS was only able categorize 55% of the empty pixels, which results in high GOFR scores for rural districts of Bioko, but when normalization for each district by population was applied, the GOFR score improved in Riaba and Baney, which are largely rural districts. Since LS is not currently available at the 100 m resolution, we do not know if it would improve performance at categorizing empty space as we observed in the HRSL and WP. The distribution of the mapped population density compared to the BIMEP population density was the most linear relationship and did not show the binning of population which is seen in the other gridded surfaces. The mis-allocated pixels on the y-axis were also distributed from both high and low population distributions without an obvious skew. LS outperforms WP at all population densities (Figure 6) and outperforms HRSL at high population densities in precision and recall. The construction of LS datasets incorporates population area weights based on administrative areas as well as land use classification down to the 1 km pixels. In the absence of reliable official local geo-referencing, this could result in the LS surface distributing the entire district population according to only the population likelihood locations and not from imagery and census data (19). Fuzzy spatial characterization of land use assignments, which is common in Africa at the fine scale gridded resolutions we are looking at, could result in an output reflective of a residential only population distribution rather than an ambient population distribution with mixed use or areas where people do not live.

### High-resolution Settlement Layer

The HRSL had the best performance overall. The HRSL still underestimated the maximum population density compared to the BIMEP gridded census data by around 66% in both the 1 km and 100 m pixel grid surfaces (Table 1). Both BIMEP and HRSL were very close in percent urban, percent empty, and overall island population total. It also provides the highest population accuracy, recall, and precision across the majority of the population density categories, especially for all the empty pixels (Table 3). The HRSL had the lowest GOFR ratio after normalization across all four districts and visually was the most like BIMEP (Figure 2). The HRSL scatter plot at 1 km had the best *R*^2^ value of any surface at 65%. Interestingly, for 100 m the HRSL scatter plot showed population density binning lines like we observed in WP, but with significantly more, and a lower *R*^2^ value of 56%, which would infer that the 1 km surface better matched the BIMEP distribution and had a more natural spread. We found the HRSL did not match the BIMEP distribution only at population densities greater than 1,000 (Figure 5B), and this was true for both 1 km and 100 m maps. The HRSL defines an urban area as 10,000 people or greater, so it is possible that the HRSL is unable to assign greater than 10,000 people to a single gridded pixel. The proportion of population by density for HRSL was closest to BIMEP at each density category (Figure 6), but while HRSL had the greatest accuracy across all population densities, there was a drop off in recall and precision at the same point before 10,000 people per pixel as before. The HRSL settlement layer most closely matched the BIMEP surface, which indicated that for Bioko Island it was the most correct human population map we examined. The challenges the HRSL population surface had characterizing high density populations bear further examination but may be due to the structural image mining approach the model is based on. Even with this consideration, our findings show that approaches to human population maps that rely more on remote sensing and image processing are better able to discern where there are not any people, which, in combination with official census data, produces an informative map.

### Malaria Burden Estimation

The disparity in the results for *Pf*PR values computed from the population density surfaces was consistent with previous studies examining the effect of population layers on malaria modeling estimates (20). The *Pf*PR estimates were very similar between WP, LS, and the HRSL but there was a noticeable difference between the estimated population fraction infected between layers when the *Pf*PR was between 10% and 30% (Figure 7A). LS tended to overestimate the population fraction at *Pf*PR around 10% compared to BIMEP but at 20% and greater had similar cumulative and probability density curves. WP had a consistently lower population fraction at *Pf*PR levels than BIMEP and along with the HRSL, however the HRSL had a smaller difference in population fraction in both the cumulative distribution and probability density functions (Figure 7B). Although the HRSL had a more similar dispersal overall compared with BIMEP, the LS surface had a more accurate distribution of population in central Malabo, which would explain its better fit to BIMEP population densities at *Pf*PR cutoffs above 15%. While the population fractions were only several percentage points different between surfaces, on an island wide scale this represents several thousand potential malaria cases. Additionally, the range where we see the difference in results is around the mesoendemic to hypoendemic transmission threshold, which is where Bioko Island’s parasite prevalence rate is currently estimated (21). Our results suggest the disparity in models is most apparent at these transition points between endemicity levels, which demonstrates the importance of using the most correct human population maps for modeling and estimating malaria.

## Conclusion

Having gold standard data, even for relatively small places such as Bioko Island, is useful as a benchmark for gridded human population density surfaces. This data provides the scale with which to evaluate the GOFR. It provides true values for accuracy profiles, whose recall metric was a strong discriminator of these maps. Even in the absence of gold standard data, plots of eCDFs for some representative area would give a detailed understanding of biases among available population maps. All of these metrics are demonstrated in the repository of code for this article, provided for Guidelines for Accurate and Transparent Health Estimates Reporting (GATHER) compliance (22, 23).

Fig. 2 gives an immediate sense of how these population maps compare, but it’s a comparison of maps available at the time of writing. New ancillary data and new algorithms may arrive this year to produce better versions of all three population maps. We are confident about the comparison because we have gold standard data for one small region and because the metrics quantify that comparison.

All of these maps are, themselves, models, which carry traces of their chosen source data and algorithms. This quantitative analysis highlighted strengths and limitations of those models, from caps on population per pixel to remarkably good identification of rural house locations. If, in the future, the HRSL could improve its estimation of the relative size of each household, it could provide both a single source for both urban and rural populations at 30 m resolution. Meanwhile, some combination of LS and HRSL would be the closest match in BIMEP.

## ACKNOWLEDGEMENTS

This work was supported by the Bill and Melinda Gates Foundation grant OPP1110495. We would like to acknowledge the BIMEP mapping team for their work collecting and processing the BIMEP population data. These data would not have been possible to realize without the support of the National Malaria Control Program and the Ministry of Health and Social Welfare of Equatorial Guinea, as well as Marathon Oil, Noble Energy, AMPCO (Atlantic Methanol Production Company) and the Ministry of Mines and Energy of Equatorial Guinea.

